# The role of reading experience in atypical cortical tracking of speech and speech-in-noise in dyslexia

**DOI:** 10.1101/2021.07.13.452155

**Authors:** Florian Destoky, Julie Bertels, Maxime Niesen, Vincent Wens, Marc Vander Ghinst, Antonin Rovai, Nicola Trotta, Marie Lallier, Xavier De Tiège, Mathieu Bourguignon

## Abstract

Dyslexia is a frequent developmental disorder in which reading acquisition is delayed and that is usually associated with difficulties understanding speech in noise. At the neuronal level, children with dyslexia were reported to display abnormal cortical tracking of speech (CTS) at phrasal rate. Here, we aimed to determine if abnormal tracking is a cause or a consequence of dyslexia and if it is modulated by the severity of dyslexia or the presence of acoustic noise.

We included 26 school-age children with dyslexia, 26 age-matched controls and 26 reading-level matched controls. All were native French speakers. Children’s brain activity was recorded with magnetoencephalography while they listened to continuous speech in noiseless and multiple noise conditions. CTS values were compared between groups, conditions and hemispheres, and also within groups, between children with best and worse reading performance.

Syllabic CTS was significantly reduced in the right superior temporal gyrus in children with dyslexia compared with controls matched for age but not for reading level. Among children with dyslexia, phrasal CTS tended to lateralize to the left hemisphere in severe dyslexia and lateralized to the right hemisphere in children with mild dyslexia and in all control groups. Finally, phrasal CTS was lower in children with dyslexia compared with age-matched controls, but only in informational noise conditions. No such effect was seen in comparison with reading-level matched controls.

Overall, our results confirmed the finding of altered neuronal basis of speech perception in noiseless and babble noise conditions in dyslexia compared with age-matched peers. However, the absence of alteration in comparison with reading-level matched controls suggests that such alterations are a consequence of reduced reading experience rather than a cause of dyslexia.

**Highlights:**

- The cortical tracking of speech (CTS) was assessed in dyslexia and proper controls
- Phrasal CTS and its resistance to noise were altered in dyslexia
- Such alterations were not found in comparison with controls matched for reading level
- The severity of dyslexia modulated the hemispheric lateralization of phrasal CTS

## 1. Introduction

Dyslexia is a developmental disorder in which reading acquisition is specifically delayed despite normal intelligence, peripheral vision and audition, appropriate schooling, and the absence of psychiatric disorders (Lyon et al., 2003). In most of the children, dyslexia would stem from a phonological deficit (Goswami, 2015; Saksida et al., 2016). Accordingly, to better understand the neural underpinnings of this deficit, many studies have sought and found traces of altered neural activity in dyslexia in tasks involving phonological processing (Bonte et al., 2007; Hämäläinen et al., 2015; Leppänen et al., 2012; Paz-Alonso et al., 2018). However, some interventional studies did not find any specific benefit of a phonological awareness training for children at high risk of dyslexia (Krashen, 1999; Olson et al., 1997; Pape-Neumann et al., 2015). Hence, phonological awareness skills correlate with upcoming reading abilities but do not determine them. So what causes both reading and phonological disorders in dyslexia?

According to the temporal sampling framework for developmental dyslexia (Goswami, 2011), abnormal temporal sampling of speech by auditory cortical oscillations would cause a deficit in both reading acquisition and phonological awareness. A tangible manifestation of the abnormal sampling would be an abnormal alignment of cortical oscillations to the different linguistic structures of speech, which can be derived from electrophysiological recordings. Indeed, when listening to connected speech, human auditory cortical activity tracks the fluctuations of speech temporal envelope at frequencies matching the occurrence rate of words/phrases/sentences (below 2 Hz) and syllables (2–8 Hz) (Ahissar et al., 2001; Bourguignon et al., 2013; Destoky et al., 2019a; Gross et al., 2013; Luo and Poeppel, 2007; Meyer et al., 2017; Meyer and Gumbert, 2018; Molinaro et al., 2016a; Vander Ghinst et al., 2019a). Such cortical tracking of speech (CTS) is thought to be essential for speech comprehension (Ahissar et al., 2001; Ding et al., 2016; Luo and Poeppel, 2007; Meyer et al., 2017; Peelle et al., 2013a; Riecke et al., 2018; Vanthornhout et al., 2018). CTS would subserve the segmentation or parsing of incoming speech for further speech recognition (Ahissar et al., 2001; Ding et al., 2016; Ding and Simon, 2014; Gross et al., 2013; Meyer et al., 2017). In line with the temporal sampling deficit hypothesis, CTS at low frequencies was found to be altered in dyslexia (Di Liberto et al., 2018a; Molinaro et al., 2016a; Power et al., 2016a). Indeed, compared with typical readers of the same age, children with dyslexia show reduced CTS at 0.5–1 Hz in both the right auditory cortex and the left inferior frontal gyrus, and reduced feedforward coupling between these two brain areas (Molinaro et al., 2016a).

However, a recent replication study did not find any CTS alteration in dyslexia (Lizarazu et al., 2021a). There are several reasons that may explain the discrepancy with the aforementioned studies. The degree of CTS alteration in dyslexia may indeed depend on (i) the language, (ii) the difficulty of the listening task, and (iii) the severity of the reading deficit present in the selected sample of dyslexic readers. Concerning the language, altered CTS was found in English and Spanish dyslexic child readers (Di Liberto et al., 2018a; Molinaro et al., 2016a; Power et al., 2016a), but not in French (Lizarazu et al., 2021a). However, in French, the lexical stress is totally predictable as it always falls on the last syllable. In contrast, lexical stress in Spanish and English changes depending on the word itself and is used to differentiate between words made of the exact same sequence of phonemes. The perfect predictability of lexical stress in French leads to a “stress deafness” in native French speakers (Dupoux et al., 1997). Because of this “stress deafness” in French, the atypical right-hemisphere neural oscillatory sampling for the low frequencies seen in English and Spanish dyslexic readers might be less severe in French dyslexic readers (Lallier et al., 2017). Concerning the difficulty of the listening task, all studies assessing CTS in dyslexia were conducted in noiseless conditions. Yet, the speech perception deficit in dyslexia is exacerbated in adverse listening conditions (Lachmann and Weis, 2018; Ziegler et al., 2009). This speech in noise (SiN) perception deficit is not due to poor spectro-temporal, low-level auditory resolution but rather to inaccurate speech representation (Lachmann and Weis, 2018), especially when the background noise is composed of speech (Calcus et al., 2015; Lachmann and Weis, 2018; Ziegler et al., 2009). Hence, the CTS alteration in dyslexia should be most salient in SiN conditions, even for French speakers. Finally, concerning the severity of the reading deficit of the included children with dyslexia, the phonological deficit is more commonly seen in severe than mild dyslexia (Saksida et al., 2016). Also along this line, reading abilities correlate with some aspects of CTS in noisy conditions (Destoky et al., 2020). Yet, none of the previous studies assessing CTS in dyslexia (Di Liberto et al., 2018a; Lizarazu et al., 2021a; Molinaro et al., 2016a; Power et al., 2016a) considered the possibility that CTS is altered only in the most severe form of dyslexia. Also, two of the three studies reporting a CTS deficit in dyslexia did not include a control group matched for reading level (Leong and Goswami, 2014; Molinaro et al., 2016a). However, it is well established that reading acquisition itself influences cognitive and cerebral functions (Carreiras et al., 2009; Goswami, 2015). One way to control for the effect of reading experience, and get novel insights into the causal link between CTS deficit and dyslexia, is to compare children with dyslexia with controls matched for the reading level in addition to the classical comparison with age-matched controls.

This study therefore aimed at determining if altered CTS is a potential cause of dyslexia. As most innovative aspects, we (i) included comparison with both controls matched for age and younger controls matched for the reading level, (ii) assessed the impact of the severity of the reading deficit on uncovered CTS alterations, and (iii) included challenging listening conditions to exacerbate potential CTS alterations in native French readers. This design was selected to answer the three following research questions: (i) Is CTS altered in French speaking children with dyslexia as a cause or a consequence of reduced reading experience? (ii) Is such alteration dependent on the severity of dyslexia? And (iii) is CTS alteration in dyslexia more salient in challenging noisy conditions, possibly depending on the severity of the reading deficit?

## 2. Methods

### 2.1. Participants

Seventy-eight children enrolled in elementary school were included in this study: 26 children with a formal diagnosis of dyslexia (*Dys*; mean ± SD age, 10.2 ± 1.1 years; 17 females), 26 typical readers matched for age (*Ctrl-Age*; 10.0 ± 1.0 years; 13 females), and 26 younger children matched for reading level (*Ctrl-Read*; 7.8 ± 0.6; 11 females). A previous study of our group has already reported on this sample, and on the outcome of the comprehensive neuropsychological evaluation they underwent (Destoky et al., 2020).

Table 1 presents the scores on which participants were matched: age, IQ, socio-economic status, and reading abilities, the latter being assessed by reading speed on lists of regular words, irregular words and pseudowords (ODEDYS-2 (Jacquier-Roux et al., 2002)), and by reading speed and accuracy on a connected text (Alouette-R test (Lefavrais, 2005)). Table 1 shows that all groups had similar IQ and socio-economic status, that dyslexic readers compared with age-matched controls had about the same age and lower reading scores, and that dyslexic readers compared with reading-level-matched controls were older and had similar reading scores. A previous analysis of the reading scores also revealed that our dyslexic readers had a rather homogenous reading profile, characterized by similar reading difficulties in the two reading pathways (Destoky et al. 2020).

**Table 1.** Mean and standard deviation of behavioral scores in each reading group of 26 children and comparisons (*t*-tests) between groups. The number of degrees of freedom was 50 for all comparisons except for socio-economic status for which some data were missing (49 for dyslexic readers vs. age-matched controls; 47 for dyslexic readers vs. reading-level-matched controls). IQ, intelligence quotient; SD, standard deviation.

|  | Dyslexic readers |  | Age-matched control |  | Reading-level-matched control |  | Dyslexic readers compared with controls |  |  |  |
| --- | --- | --- | --- | --- | --- | --- | --- | --- | --- | --- |
|  | Mean | SD | Mean | SD | Mean | SD | age |  | reading level |  |
|  |  |  |  |  |  |  | *p* | *t* | *p* | *t* |
| Chronological age | 10.2 | 1.08 | 9.97 | 1.01 | 7.76 | 0.60 | 0.36 | 0.93 | **<0.0001** | **10.3** |
| Non-verbal IQ | 111 | 11 | 114 | 10 | 112 | 9 | 0.30 | -1.04 | 0.784 | -0.28 |
| Socio-economic status | 6.12 | 2.44 | 6.96 | 1.45 | 6.96 | 2.47 | 0.14 | -1.50 | 0.17 | -1.40 |
| Alouette reading accuracy | 89.0 | 5.7 | 96.2 | 2.1 | 89.0 | 6.46 | **<0.0001** | **-6.07** | 0.988 | 0.01 |
| Alouette reading speed | 141 | 61 | 292 | 91 | 138 | 64 | **<0.0001** | **-7.04** | 0.867 | 0.17 |
| Irregular words reading [words/s] | 0.54 | 0.33 | 1.16 | 0.44 | 0.40 | 0.35 | **<0.0001** | **-5.82** | 0.15 | 1.47 |
| Regular words reading [words/s] | 0.73 | 0.41 | 1.35 | 0.41 | 0.61 | 0.35 | **<0.0001** | **-5.51** | 0.29 | 1.06 |
| Pseudo-words reading [words/s] | 0.42 | 0.24 | 0.78 | 0.30 | 0.39 | 0.21 | **<0.0001** | **-4.88** | 0.61 | 0.50 |

All children were native French speakers, reported being right-handed, had normal hearing according to pure-tone audiometry (normal hearing thresholds between 0–25 dB HL for 250, 500, 1,000, 2,000, 4,000, and 8,000 Hz) and normal SiN perception as revealed by a SiN test (Lafon 30) from a French language central auditory battery (Demanez et al., 2003).

This study was approved by the local ethics committee (Comité d’Ethique Hospitalo-Facultaire Erasme-ULB, 021/406, Brussels, Belgium; approval number: P2017/081) and conducted according to the principles expressed in the Declaration of Helsinki. Participants were recruited mainly from local schools through flyer advertisements or from social networks. Participants and their legal representatives signed a written informed consent before participation. Participants were compensated with a gift card worth 50 euros.

### 2.2. Reading subgroups

As a preliminary step to test our working hypothesis that CTS is modulated by the severity of the reading deficit in dyslexia, we partitioned the *Dys* group into two subgroups maximally differing on their reading abilities. To do so, the 5 reading scores (see Table 1) were first corrected for age, time spent at school and IQ as done in our previous study (Destoky et al., 2020), and further standardized. We then used the k-means clustering algorithm implemented in MATLAB to identify the 2 subgroups. Since all the reading scores in one subgroup were higher than those in the second subgroup, we refer to them as the mild (*Dys-Mild*; *n* = 16; 11 females; mean ± SD age, 10.4 ± 1.0 years) and severe subgroups (*Dys-Severe*; *n* = 10; 6 females; 10.0 ± 1.2 years). Obviously, the 2 subgroups displayed significant differences in reading skills (Table 2). Most importantly, they did not differ significantly on their age (*t*_24_ = 0.86, *p* = 0.40) or socio-economic level (*t*_23_ = -0.13, *p* = 0.90).

**Table 2.** Mean of the standardized reading scores (i.e., z-scores) for the *Dys-Mild* and *Dys-Severe* subgroups and significance of their comparisons. Scores were standardized within the *Dys* group.

| | *Dys-Mild*<br>($n = 16$) | *Dys-Severe*<br>($n = 10$) | *p* |
| --- | --- | --- | --- |
| Text (Alouette) reading accuracy | 0.36 | -0.58 | 0.015 |
| Text (Alouette) reading speed | 0.60 | -0.96 | < 0.0001 |
| Irregular word reading speed | 0.62 | -1.00 | < 0.0001 |
| Regular word reading speed | 0.67 | -1.06 | < 0.0001 |
| Pseudo-word reading speed | 0.59 | -0.95 | < 0.0001 |

The same procedure was used to partition each of the two control groups. The *Ctrl-Age* group was split into one subgroup with high reading scores (*Ctrl-Age-High*; *n* = 12; 5 females; 10.2 ± 1.1 years) and one with low reading scores (*Ctrl-Age-Low*; *n* = 14; 8 females; 9.8 ± 0.9 years), which again did not differ on age (*t*_24_ = 0.96, *p* = 0.34) or socioeconomic level (*t*_24_ = 1.83, *p* = 0.08). Likewise, the *Ctrl-Read* group was split into a subgroup with high reading scores (*Ctrl-Read-High*; *n* = 12; 6 females; 7.82 ± 0.56) and a subgroup with low reading scores (*Ctrl-Read-Low*; *n* = 14; 7 females; 7.71 ± 0.64) not differing on age (*t*_24_ = 0.45, *p* = 0.66) or socioeconomic level (*t*_22_ = 0.60, p = 0.55). Importantly, three sets of comparisons demonstrated that each of the control subgroups remained a good control for its corresponding *Dys* subgroup: (*i*) there was no significant difference in age between *Dys-Mild* and *Ctrl-Age-High* (*t*_26_ = 0.51, *p* = 0.61) nor between *Dys-Severe* and *Ctrl-Age-Low* (*t*_26_ = 0.50, *p* = 0.62), (*ii*) there was no significant difference in reading scores between *Dys-Mild* and *Ctrl-Read-High* (*p* > 0.17 in all 5 comparisons) nor between the *Dys-Severe* and *Ctrl-Read-Low* (*p* > 0.21 in all 5 comparisons), and (*iii*) there was no significant difference in socioeconomic level between *Dys-Mild* and the two control groups with high reading scores (*Ctrl-Age-High*, *t*_26_ = -1.89, *p* = 0.070; *Ctrl-Read-High*, *t*_26_ = -1.67, *p* = 0.11) nor between *Dys-Severe* and the two control groups with low reading scores (*Ctrl-Age-Low*, *t*_22_ = -0.24, *p* = 0.82; *Ctrl-Read-Low*, *t*_22_ = -0.22, *p* = 0.83).

### 2.3. Stimuli

Figure 1 illustrates the time-course of the video stimuli, which were exactly the same as in a previous study from our group (for more details, see Destoky et al. 2020). Video stimuli were derived from 12 audiovisual recordings of four native French speaking narrators (two females, three recordings per narrator) telling a story for approximately 6 min (mean ± SD, 6.0 ± 0.8 min). In each video, the first 5 s were kept unaltered to enable children to unambiguously identify the narrator’s voice and face that they were requested to attend to. The remainder of the video was divided into 10 consecutive blocks of equal size that were assigned to nine conditions. Two blocks were assigned to the noiseless condition, in which the audio track was kept unaltered but the video was replaced by static pictures illustrating the story (mean ± SD picture presentation time across all videos, 27.7 ± 10.8 s). The remaining eight blocks were assigned to eight conditions in which the original sound was mixed with a background noise at 3 dB signal-to-noise ratio. There were four different types of noise, and each type of noise was presented once with the original video, thereby giving access to visual speech information, and once with the static pictures illustrating the story and hence without visual speech information. Here, as the aim was to use the most challenging listening condition, we report only on the data in which visual speech information was absent since it is already well documented that phrasal and syllabic CTS in noise is boosted when visual speech information is available (Destoky et al., 2020; Golumbic et al., 2013; Park et al., 2018, 2016). The different types of noise differed in the degree of energetic and informational interference they introduced (Pollack, 1975). Nevertheless, in our previous study on the same data (Destoky et al., 2020), we observed that the degree of energetic masking had little impact on CTS values. For this reason, we pooled the data and considered only the distinction between non-speech (non-informational) noises and babble (informational) noises. The non-speech noises were a white noise high-pass filtered at 10,000 Hz or a noise spectro-temporally matched to the narrator’s voice. The babble noises were five-talker cocktail party noises of the same gender as the narrator or of the opposite gender. Individual noise components were obtained from a French audiobook database (http://www.litteratureaudio.com), normalized, and mixed linearly. The assignment of conditions to blocks was random. Ensuing videos were grouped into three disjoint sets featuring one video per narrator (total set duration: 23.0, 24.3, 24.65 min), and there were four versions of each set differing in condition random ordering.

**Figure 1.**
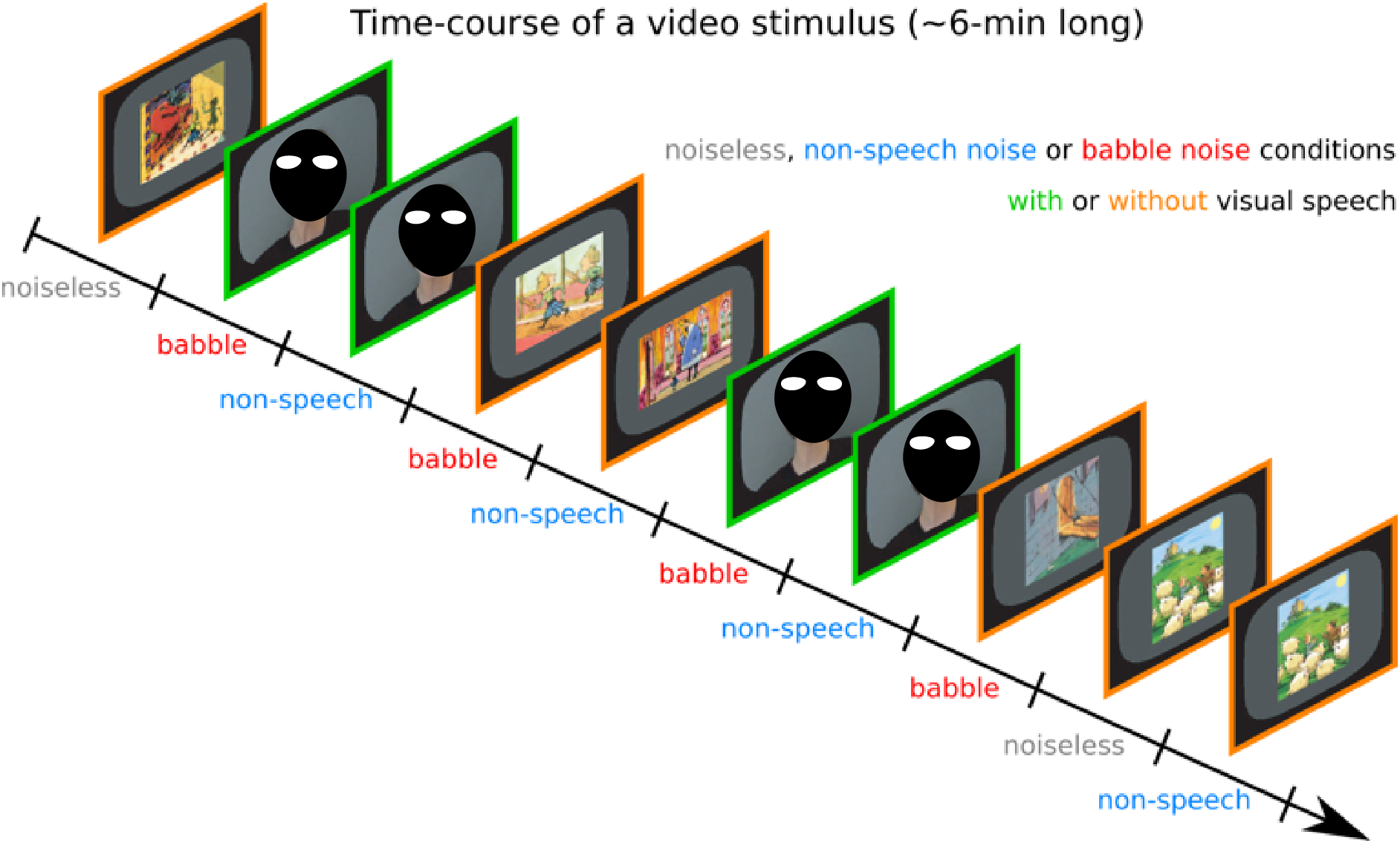
Illustration of the time-course of a video stimulus.

### 2.4. Experimental paradigm

Participants laid on a bed with their head inside a magnetoencephalography (MEG) helmet. Their brain activity was recorded while they were attending four videos of a randomly selected set presented in a random order (separate recording for each video), and finally while they were at rest (eyes opened, fixation cross) for 5 min. They were instructed to watch the videos attentively, listen to the narrators’ voice while ignoring the interfering noise, and remain as still as possible. After each video, they were asked 10 yes/no simple comprehension questions. Videos were projected onto a back-projection screen placed vertically, approximately 120 cm away from the MEG helmet. The inner dimensions of the black frame were 35.2 cm (horizontal) and 28.8 cm (vertical), and the narrator’s face spanned approximately 15 cm (horizontal) and approximately 20 cm (vertical). Participants could see the screen through a mirror placed above their head. In total, the optical path from the screen to participants’ eyes was approximately 150 cm. Sounds were delivered at 60 dB (measured at ear level) through a MEG-compatible, front-facing, flat-panel loudspeaker (Panphonics Oy, Espoo, Finland) placed approximately 1 m behind the screen.

### 2.5. Data acquisition

During the experimental conditions, participants’ brain activity was recorded with MEG at the CUB Hôpital Erasme (Brussels, Belgium). Neuromagnetic signals were recorded with a whole-scalp–covering MEG system (Triux, MEGIN, Croton Healthcare, Helsinki, Finland) placed in a lightweight, magnetically shielded room (Maxshield, MEGIN, Croton Healthcare, Helsinki, Finland), the characteristics of which have been described elsewhere (De Tiège et al., 2008). The sensor array of the MEG system comprised 306 sensors arranged in 102 triplets of one magnetometer and two orthogonal planar gradiometers. Magnetometers measure the radial component of the magnetic field, whereas planar gradiometers measure its spatial derivative in the tangential directions. MEG signals were band-pass filtered at 0.1–330 Hz and sampled at 1,000 Hz.

We used four head-position indicator coils to monitor the subjects’ head position during the experimentation. Before the MEG session, we digitized the location of these coils and at least 300 head-surface points (on scalp, nose, and face) with respect to anatomical fiducials with an electromagnetic tracker (Fastrack, Polhemus).

Finally, subjects’ high-resolution 3D T1-weighted cerebral images were acquired with a 3T hybrid PET-MR scanner (SIGNA, GE Healthcare, Milwaukee, Wisconsin, USA) after the MEG session.

### 2.6. Data preprocessing

Continuous MEG data were first preprocessed off-line using the temporal signal space separation method implemented in MaxFilter software (MaxFilter, MEGIN; correlation limit 0.9, segment length 20 s) to suppress external interferences and to correct for head movements (Taulu et al., 2005; Taulu and Simola, 2006). To further suppress physiological artifacts, 30 independent components were evaluated from the data band-pass filtered at 0.1–25 Hz and reduced to a rank of 30 with principal component analysis. Independent components corresponding to heartbeat, eye-blink, and eye-movement artifacts were identified, and corresponding MEG signals reconstructed by means of the mixing matrix were subtracted from the full-rank data. An ANOVA revealed no significant difference between groups (*F*_2,50_ = 1.03, *p* = 0.36) in the number of subtracted independent components (mean ± SD; *Dys*, 3.3 ± 0.6; *Ctrl-Age*, 3.6 ± 0.9; *Ctrl-Read*, 3.3 ± 0.6). Finally, time points at timings 1 s around remaining artifacts were set to bad. Data were considered contaminated by artifacts when MEG amplitude exceeded 5 pT in at least one magnetometer or 1 pT/cm in at least one gradiometer.

We extracted the temporal envelope of the attended speech (i.e., narrators’ voice) using a state-of-the-art approach (Biesmans et al., 2017). Briefly, audio signals were band-pass filtered using a gammatone filter bank (15 filters centered on logarithmically spaced frequencies from 150 Hz to 4,000 Hz), and sub-band envelopes were computed using Hilbert transform, elevated to the power 0.6, and averaged across bands.

### 2.7. CTS estimated globally for the left and right hemispheres

For each condition and participant, a global value of cortical tracking of the attended speech was evaluated for all left-hemisphere sensors at once and for all right-hemisphere sensors at once. Using the mTRF toolbox (Crosse et al., 2016), we trained a decoder on MEG data to reconstruct speech temporal envelope and estimated its Pearson correlation with real speech temporal envelope. This correlation is often referred to as the reconstruction accuracy, and it provides a global measure of CTS. In brief, electrophysiological data were band-pass filtered at 0.2–1.5 Hz (phrasal rate) or 2–8 Hz (syllabic rate) and a decoder for speech temporal envelope was built based on MEG data from –500 ms to 1000 ms (phrasal) or from 0 ms to 250 ms (syllabic) with respect to speech temporal envelope. The decoder used to estimate reconstruction accuracy in a given condition was built based on the data in all the other conditions, using 10-fold cross-validation to select the optimal regularisation applied to limit the norm of the derivative of the reconstructed speech temporal envelope (Crosse et al., 2016). For a full description of the procedure, see our previous study (Destoky et al., 2020).

### 2.8. CTS estimated in the source space

As a preliminary step to estimate brain maps of CTS, MEG signals were projected into the source space. For that, MEG and MRI coordinate systems were co-registered using the 3 anatomical fiducial points for initial estimation and the head-surface points for further manual refinement. When a participant’s MRI was missing (n = 27), we used that of another participant of roughly the same age, which we linearly deformed to best match head-surface points using the CPD toolbox (Myronenko and Song, 2010) embedded in FieldTrip (Donders Institute for Brain Cognition and Behaviour, Nijmegen, The Netherlands, RRID:SCR_004849; (Oostenveld et al., 2011)). The individual MRIs were segmented using the Freesurfer software (Martinos Center for Biomedical Imaging, Boston, MA, RRID:SCR_001847; (Reuter et al., 2012)). Then, a non-linear transformation from individual MRIs to the MNI brain was computed using the spatial normalization algorithm implemented in Statistical Parametric Mapping (SPM8, Wellcome Department of Cognitive Neurology, London, UK, RRID:SCR_007037; (Ashburner et al., 1997; Ashburner and Friston, 1999)). This transformation was used to map a homogeneous 5-mm grid sampling the MNI brain volume onto individual brain volumes. For each subject and grid point, the MEG forward model corresponding to three orthogonal current dipoles was computed using the one-layer Boundary Element Method implemented in the MNE software suite (Martinos Centre for Biomedical Imaging, Boston, MA, RRID:SCR_005972; (Gramfort et al., 2014)). The forward model was then reduced to its two first principal components. This procedure is justified by the insensitivity of MEG to currents radial to the skull, and hence, this dimension reduction leads to considering only the tangential sources. Source signals were then reconstructed with Minimum-Norm Estimates inverse solution (Dale and Sereno, 1993).

We followed a similar approach to that used at the sensor level to estimate source-level CTS. For each grid point, we trained a decoder on the two-dimensional source time-series to reconstruct speech temporal envelope. Again, the decoder was trained on the data from all but one condition, and used to estimate CTS in the left-out condition. To speed up computations, the training was performed without cross-validation, with the ridge value retained in a sensor-space analysis run on all gradiometer sensors at once. This procedure yielded a source map of CTS for each participant, condition, and frequency range of interest; and because the source space was defined on the MNI brain, all CTS maps were inherently corregistered with the MNI brain. Hence, group-averaged maps were simply produced as the mean of individual maps within age groups, conditions and frequency ranges of interest.

We further identified the coordinates of local maxima in group-averaged CTS maps. Such local maxima of CTS are sets of contiguous voxels displaying higher CTS values than all neighbouring voxels (Bourguignon et al., 2012). We only report statistically significant local maxima of CTS, disregarding the extents of these clusters. Indeed, cluster extent is hardly interpretable in view of the inherent smoothness of MEG source reconstruction (Bourguignon et al., 2018; Hämäläinen and Ilmoniemi, 1994; Wens et al., 2015).

We also estimated the contrast of source maps between the different groups and identified the coordinates of local maxima therein.

### 2.9. Statistical analyses

#### 2.9.1. Effect of hemisphere, group, subgroup, and noise on CTS

To test whether CTS is altered in dyslexia in comparison with controls in age or reading level, potentially in different ways in the left-versus right hemisphere (first research question), we ran a repeated measures ANOVA on CTS values in the noiseless condition with factors hemisphere and group (i.e., *Dys, Ctrl-Age* and *Ctrl-Read*), separately for phrasal and syllabic CTS.

To test whether CTS is affected by the severity of the reading deficit in dyslexia (second research question), we ran the same analysis as above, but only on the data for *Dys*, with factors hemisphere and subgroup (*Mild* vs. *Severe*), separately for phrasal and syllabic CTS.

Finally, to test whether CTS alteration in dyslexia is most visible in SiN conditions (third research question), possibly depending on the severity of the reading deficit and on the hemisphere, we assessed with a linear mixed-effects analysis implemented in R (Team and Others, 2013) and lme4 (Bates et al., 2015) the effect of hemisphere, group, subgroup (*Mild/High vs. Severe/Low*) and noise (noiseless, non-speech, babble) on CTS values, separately for phrasal and syllabic CTS. An ANOVA could not be used here, because it cannot accommodate the clustering of participants in both groups and subgroups. We followed a step-up approach to iteratively identify all statistically significant effects. In brief, we started with a null model that included only a different random intercept for each subject. The model was iteratively compared with models incremented with simple fixed effects added one by one. At every step, the most significant fixed effect was retained until the addition of the remaining effects did not improve the model any further (*p* > 0.05). The same procedure was then repeated to refine the ensuing model with the interactions of the simple fixed effects of order 2, 3 and then 4.

In all analyses, post-hoc t-tests were conducted to clarify the effects uncovered with the ANOVAs or linear mixed-effects analysis.

#### 2.9.2. Significance of local maxima of CTS

The statistical significance of CTS local maxima observed in group-averaged maps for each group (i.e., *Dys, Ctrl-Age and Ctrl-Read*) and frequency range of interest (i.e., 0.2–1.5 and 2–8 Hz) was assessed with a non-parametric permutation test that intrinsically corrects for multiple spatial comparisons (Nichols and Holmes, 2002). First, participant and group-averaged *null* maps of CTS were computed with MEG and voice signals in each story rotated in time by about half of story length (i.e., the first and second halves were swapped, thereby destroying genuine coupling but preserving spectral properties). The exact temporal rotation applied was chosen to match a pause in speech to enforce continuity. Group-averaged difference maps were obtained by subtracting *genuine* and *null* group-averaged CTS maps. Under the null hypothesis that CTS maps are the same whatever the experimental condition, the labeling *genuine* or *null* are exchangeable prior to difference map computation (Nichols and Holmes, 2002). To reject this hypothesis and to compute a significance level for the correctly labeled difference map, the sample distribution of the maximum of the difference map’s absolute value within the entire brain was computed from a subset of 1000 permutations. The threshold at *p* < 0.05 was computed as the 95^th^ percentile of the sample distribution (Nichols and Holmes, 2002). All supra-threshold local maxima of CTS were interpreted as indicative of brain regions showing statistically significant CTS and will be referred to as sources of CTS.

Permutation tests can be too conservative for voxels other than the one with the maximum observed statistic (Nichols and Holmes, 2002). For example, dominant CTS values in the right auditory cortex could bias the permutation distribution and overshadow weaker CTS values in the left auditory cortex, even if these were highly consistent across subjects. Therefore, the permutation test described above was conducted separately for left- and right-hemisphere voxels.

The same approach was used to assess the significance of local maxima in contrasts between group maps.

## 3. Results

### 3.1. Is CTS altered in French speaking children with dyslexia as a cause or consequence of reduced reading experience?

Figure 2 presents the values of phrasal/sentential and syllabic CTS in the noiseless condition for both hemispheres and the 3 groups.

**Figure 2.**
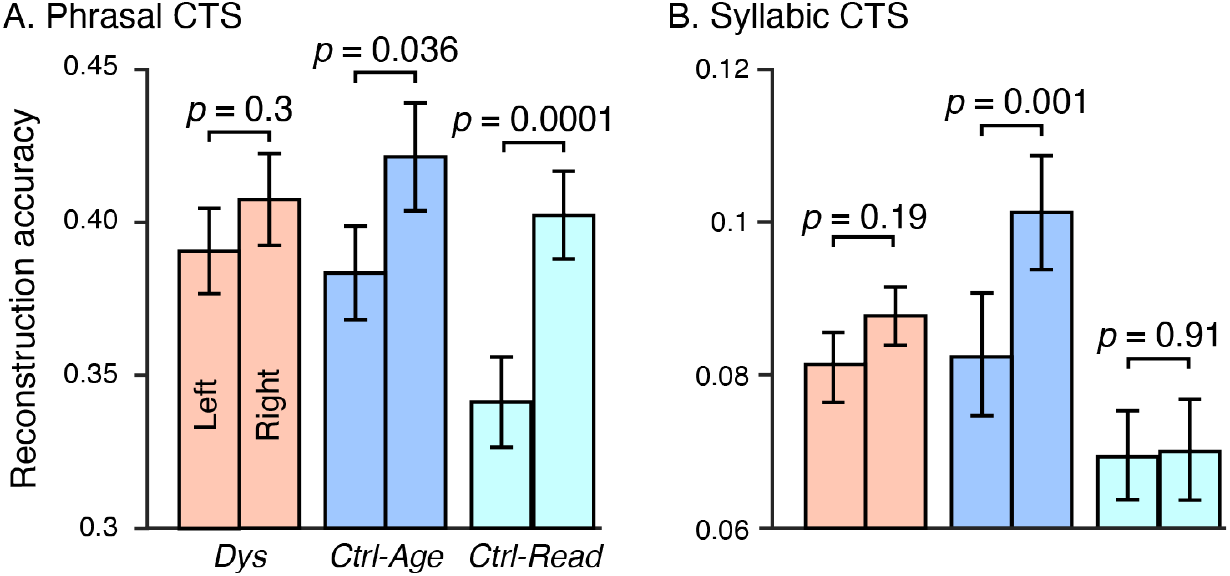
Phrasal (A) and Syllabic (B) CTS estimated with reconstruction accuracy in each hemisphere and group. Bars and vertical lines indicate mean ± SEM values. P-values are provided for comparisons between hemispheres.

The ANOVA run on these phrasal CTS values revealed a significant effect of hemisphere (*F*_1,25_ = 13.0, *p* = 0.0014), no significant effect of group (*F*_2,25_ = 1.59, *p* = 0.21) and a marginally significant interaction between hemisphere and group (*F*_2,25_ = 2.63, *p* = 0.082). Post-hoc analyses revealed that CTS was higher in the right hemisphere than in the left hemisphere in *Ctrl-Age* (*t*_25_ = 2.21, *p* = 0.036) and *Ctrl-Read* (*t*_25_ = 4.67, *p* = 0.0001) but not in *Dys* (*t*_25_ = 1.06, *p* = 0.30). Still, *Dys* and *Ctrl-Age* did not differ significantly on their level of phrasal CTS in the left hemisphere (*t*_25_ = 0.3255, *p* = 0.75) or right hemisphere (*t*_25_ = -0.78, *p* = 0.44), nor on the difference thereof (*t*_25_ = -1.06, *p* = 0.298). The latter difference between left- and right-hemisphere CTS differed between *Dys* and *Ctrl-Read* (*t*_25_ = -2.58, *p* = 0.01), and did not differ between *Ctrl-Age* and *Ctrl-Read* (*t*_25_ = -1.12, *p* = 0.27).

The ANOVA run on syllabic CTS values revealed a significant effect of hemisphere (*F*_1,25_ = 4.86, *p* = 0.037), a significant effect of group (*F*_2,25_ = 3.76, *p* = 0.030), and a significant interaction (*F*_2,25_ = 3.28, *p* = 0.046). Post-hoc analyses revealed that CTS hemispheric dominance in *Dys* and *Ctrl-Read* differed significantly from that in *Ctrl-Age* (see Fig. 2B). Indeed, the difference between right- and left-hemisphere CTS values was significantly higher in *Ctrl-Age* compared with *Ctrl-Read,* (*t*_25_ = 3.17, *p* = 0.0040), marginally higher in *Ctrl-Age* compared with *Dys* (*t*_25_ = -1.78, *p* = 0.088), and there was no significant difference between *Dys* and *Ctrl-Read* (*t*_25_ = 0.71, *p* = 0.48). Moreover, CTS was significantly higher in the right-compared with left hemisphere in *Ctrl-Age* (*t*_25_ = -3.67, *p* = 0.0011) but not in *Dys* (*t*_25_ = -1.36, *p* = 0.19) or *Ctrl-Read* (*t*_25_ = -0.11, *p* = 0.91).

In summary, phrasal CTS was not significantly altered in *Dys* compared with *Ctrl-Age*. However, syllabic CTS in *Dys* failed to show the right-hemispheric lateralization seen in *Ctrl-Age* and was akin to that in *Ctrl-Read*.

Since a global analysis that distinguishes only between left- and right-hemisphere CTS may overlook subtle differences in specific brain regions, we also compared source-space CTS maps between groups.

Significant local maxima of phrasal CTS localized in all groups in posterior superior temporal gyrus (pSTG) bilaterally, less than 5 mm from the mean coordinate across groups (left, MNI coordinates [–46 –26 2] mm; right, [58 –21 0] mm), and in the right inferior frontal gyrus, less than 9 mm from the mean coordinate across groups ([54 20 3] mm; Figure 3). The group comparisons revealed no significant difference between any of the groups in any of the hemispheres (*p* > 0.1 for the 6 comparisons).

**Figure 3.**
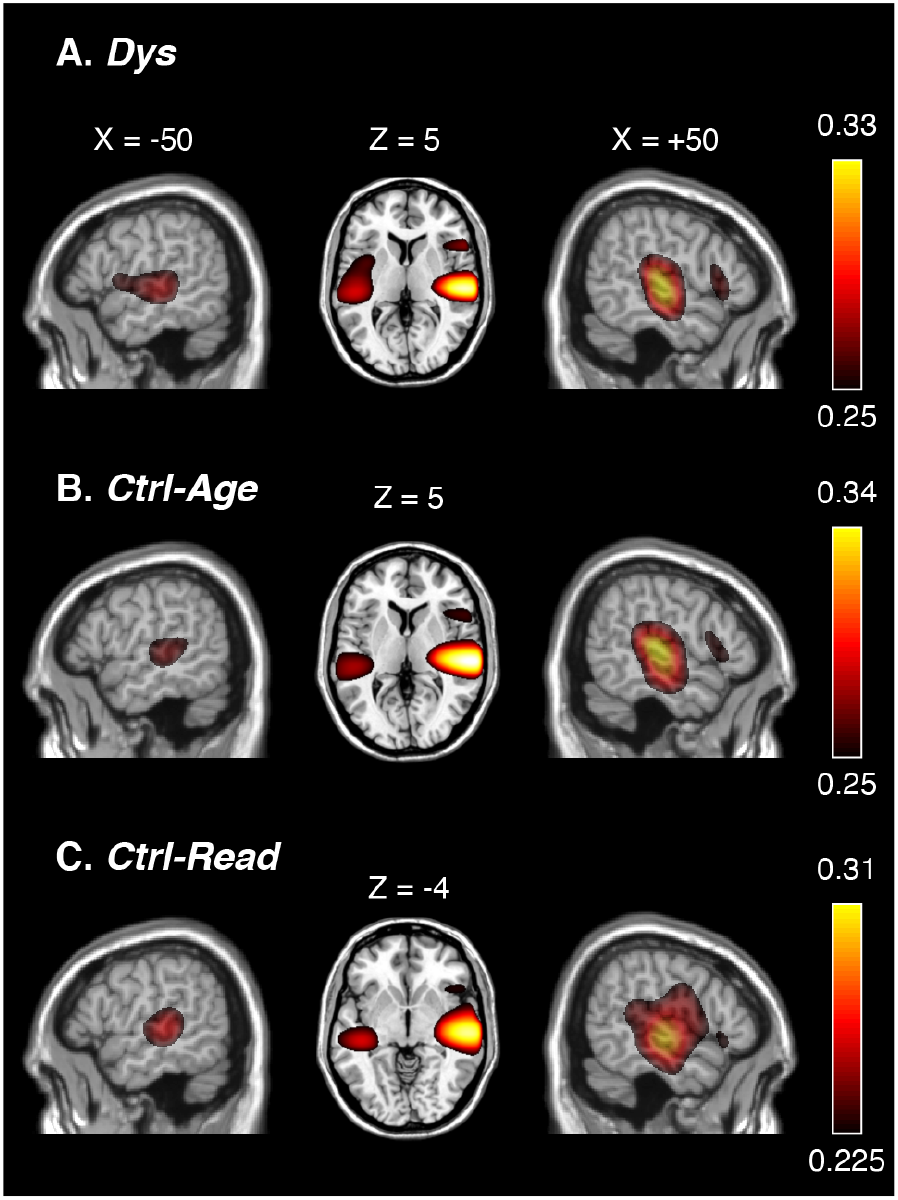
Source-level maps of phrasal CTS in the noiseless condition. A — Maps for the Dys group. B — Maps for the Ctrl-Age group. C — Maps for the Ctrl-Read group. All maps are thresholded at statistical significance level corrected for multiple comparisons across each hemisphere.

Significant local maxima of syllabic CTS localized in all groups in superior temporal gyrus (STG) bilaterally, less than 7 mm from the mean coordinate across groups (left, [–49 – 18 8] mm; right, [57 –13 8] mm) and in inferior frontal gyrus bilaterally, less than 10 mm from the mean coordinate across groups (left, [–48 20 –7] mm; right, [52 28 0] mm) (Figure 4). The group comparison revealed significantly higher CTS in the right STG in *Ctrl-Age* compared with *Dys* (*p* = 0.02 at [65 -36 -1] mm) and in *Ctrl-Age* compared with *Ctrl-Read* (*p* = 0.001 at [65 -30 10] mm), and no other significant differences (*p* > 0.07 for the 4 comparisons).

**Figure 4.**
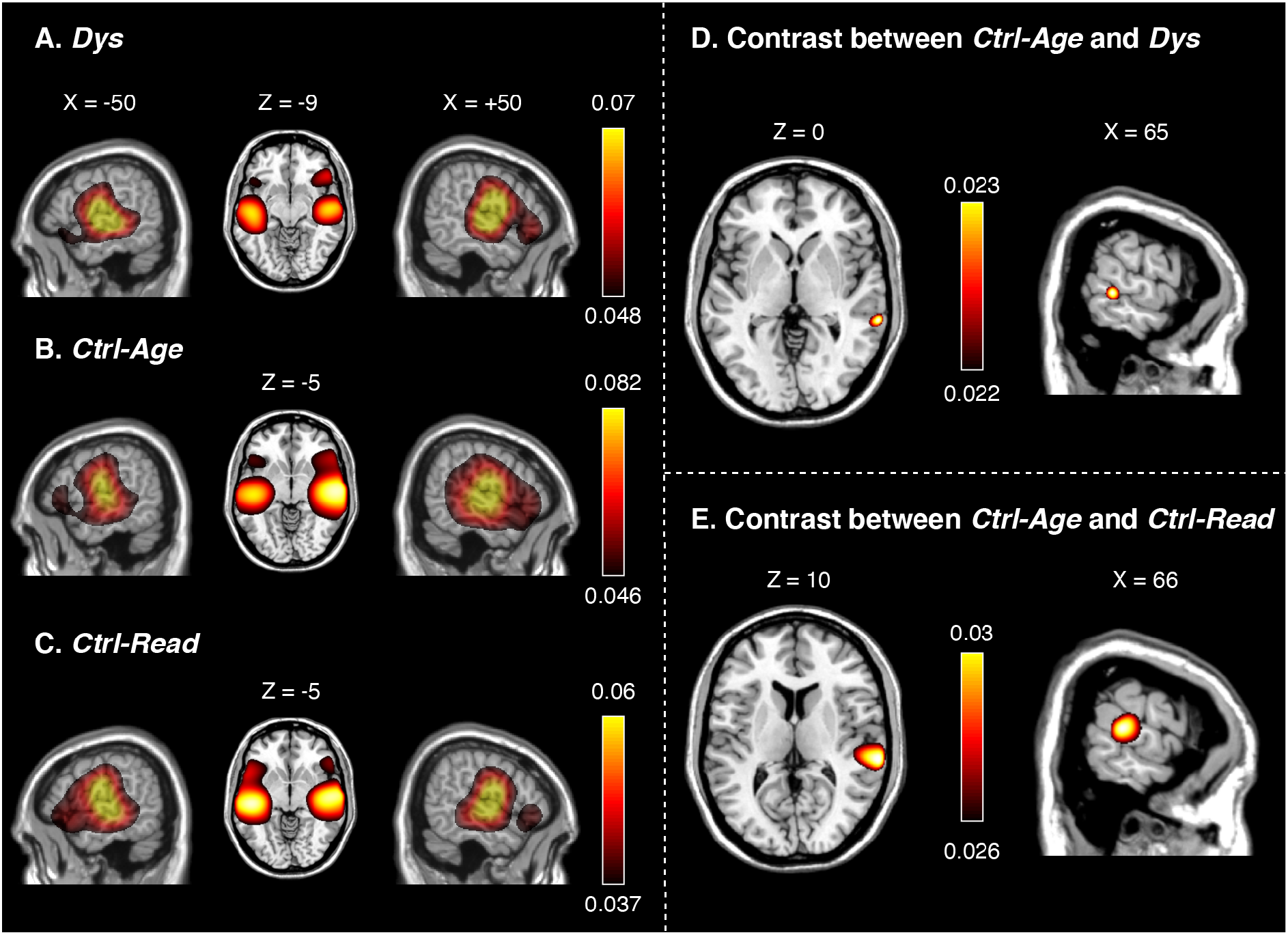
Source-level maps of syllabic CTS in the noiseless condition. A — Maps for the Dys group. B — Maps for the Ctrl-Age group. C — Maps for the Ctrl-Read group. D — Contrast between Ctrl-Age and Dys. E — Contrast between Ctrl-Age and Ctrl-Read. All maps are thresholded at statistical significance level corrected for multiple comparisons across each hemisphere.

### 3.2. Does the severity of dyslexia impact CTS?

Figure 5 presents the values of phrasal (Figure 5A) and syllabic (Figure 5B) CTS in both hemispheres and in both subgroups of dyslexic readers.

**Figure 5.**
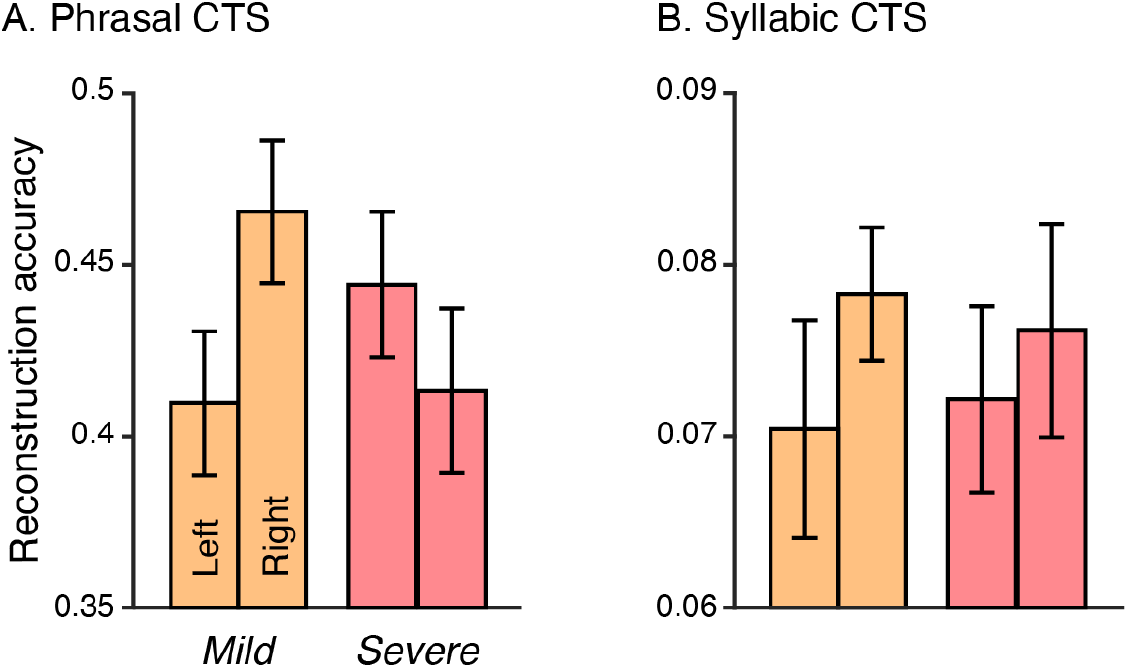
Phrasal (A) and Syllabic (B) CTS in both hemispheres for the two dyslexic subgroups (Mild and Severe). Bars and vertical lines indicate mean ± SEM values.

The ANOVA run on phrasal CTS revealed no significant effect of hemisphere (*F*_1,25_ = 0.56, *p* = 0.46), no significant effect of subgroup (*F*_1,25_ = 0.1, *p* = 0.76), but a significant interaction between hemisphere and subgroup (*F*_1,25_ = 6.79, *p* = 0.016). Post-hoc analyses revealed that CTS hemispheric dominance in *Dys-Mild* differed from that in *Dys-Severe*. Indeed, in *Dys-Mild*, CTS values were higher in the right-compared with the left hemisphere (*t*_15_ = -2.48, *p* = 0.02) while the reverse pattern was present—though not significantly—in *Dys-Severe* (*t*_9_ = 1.43, *p* = 0.19).

An ANOVA run on syllabic CTS revealed no significant effect of hemisphere (*F*_1,25_ = 1.8, *p* = 0.19), no significant effect of subgroup (*F*_1,25_ = 0.0008, *p* = 0.98), and no significant interaction between the hemisphere and subgroup (*F*_1,25_ = 0.19, *p* = 0.67).

In sum, we found that the severity of the reading deficit in dyslexia modifies the hemispheric dominance of phrasal CTS. The next subsection will clarify how these profiles compare with those in control groups.

### 3.3. Is CTS alteration in dyslexia most salient in challenging listening conditions?

Linear mixed-effects modeling of phrasal CTS values in all hemispheres, noise conditions, groups and subgroups revealed a statistically significant effect of noise (χ^2^(2) = 345, *p* < 0.0001), hemisphere (χ^2^(1) = 18.2, *p* < 0.0001), and group (χ^2^(2) = 8.75, *p* = 0.01), and significant interactions between hemisphere and noise (χ^2^(2) = 12.2, *p* = 0.0022), noise and group (χ^2^(4) = 9.61, *p* = 0.047), and hemisphere, group and subgroup (χ^2^(8) = 21.4, *p* = 0.0062).

Figure 6A illustrates the interaction between hemisphere and noise. The interaction was explained by a right hemispheric dominance for phrasal CTS in *noiseless* (*t*_77_ = -4.08, *p* = 0.0001) and *non-speech* noise conditions (*t*_77_ = -3.88, *p* = 0.0002) but not in the *babble* noise condition (*t*_77_ = 0.0.46, *p* = 0.65). A pronounced effect of babble noise was also evident, with lower phrasal CTS in babble noise condition compared with noiseless and non-speech noise conditions (*p* < 0.0001). These results replicate the finding that noise impacts more right-than left-hemisphere phrasal CTS (Destoky et al., 2020, 2019b; Vander Ghinst et al., 2019b, 2016).

**Figure 6.**
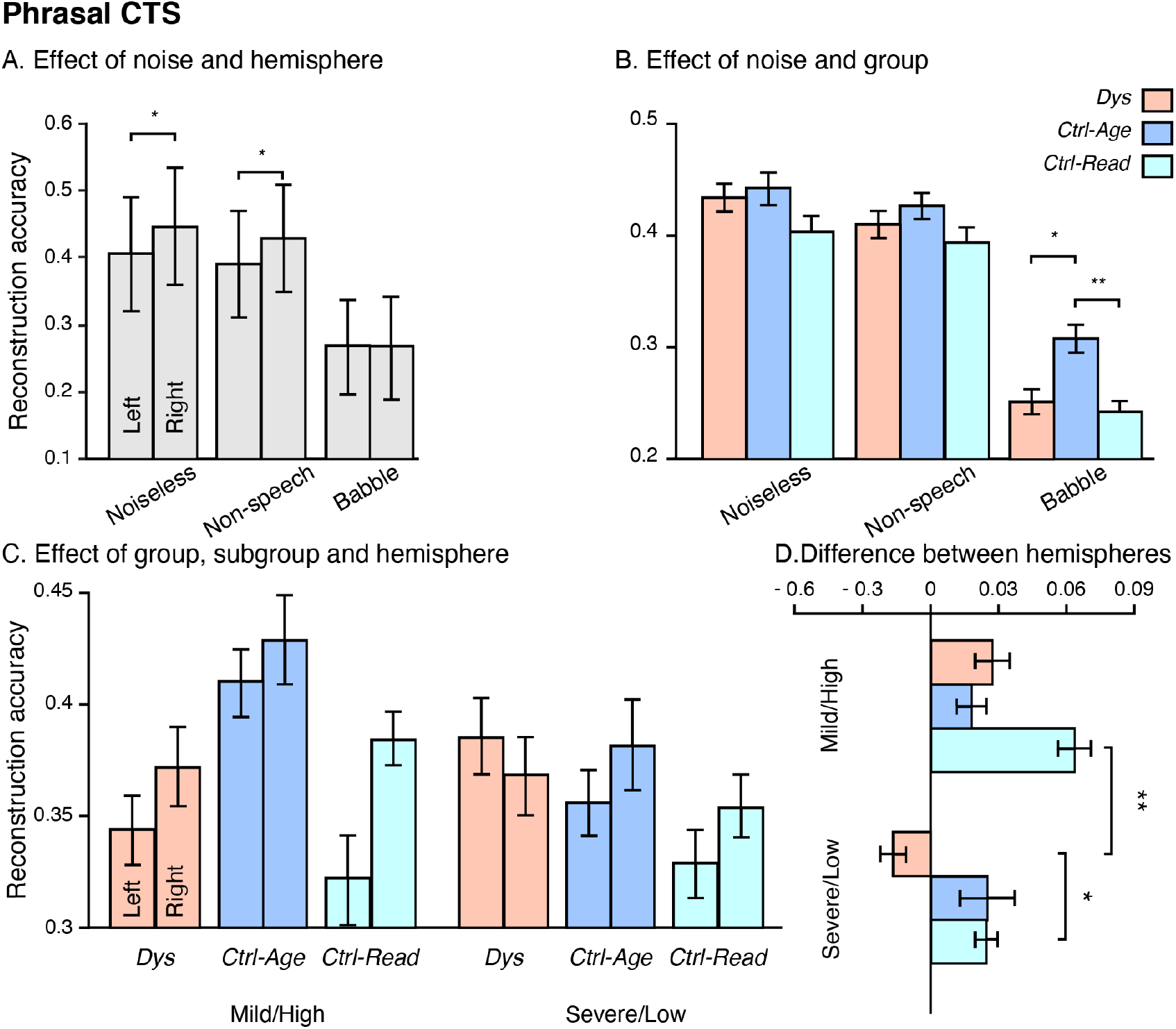
Effect of hemisphere, noise, group and subgroup on phrasal CTS. A — Phrasal CTS averaged across groups. B — Phrasal CTS averaged across hemisphere and subgroups. C — Phrasal CTS averaged across noise conditions. D — Difference between right- and left-hemisphere phrasal CTS further averaged across noise conditions. Bars and vertical (A-C) or horizontal (D) lines indicate mean ± SEM values. *, p < 0.05; **, p < 0.01.

Figure 6B illustrates the interaction between noise and group. The interaction was explained by significantly higher CTS in babble noise in *Ctrl-Age* compared with the two other groups (*Dys*, *t*_50_ = -3.11, *p* = 0.0031; *Ctrl-Read*, *t*_50_ = 3.81, *p* = 0.0004) for which there was no significant difference (*t*_50_ = 0.57, *p* = 0.57), while the groups did not differ in *noiseless* nor in *non-speech* noise conditions (*p* > 0.08 in all 6 comparisons). This indicates that the introduction of babble noise brought about a significant alteration in phrasal CTS in dyslexia and *Ctrl-Read* in comparison with *Ctrl-Age*. In other words, in reaction to babble noise, phrasal CTS in dyslexia is hindered much like in *Ctrl-Read*. Also worth noting, this effect was independent of the hemisphere and subgroup.

Figure 6C illustrates the triple interaction between hemisphere, group and subgroup. This interaction indicates that the modulation in CTS hemispheric lateralization by reading proficiency seen in *Dys* (see previous subsection) differed between groups. Importantly, this effect did not interact with noise (no quadruple interaction; χ^2^(16) = 14.21, *p* = 0.58), indicating that noise did not exacerbate the impact of reading abilities on the hemispheric dominance in *Dys*. The interaction was driven by a difference in phrasal CTS between hemispheres that differed in *Dys-Severe* and *Ctrl-Read-High* from the 4 other groups. To substantiate this effect, we compared subgroups for their difference in CTS between right and left hemispheres (see Figure 6D). As a result, the difference in *Dys-Severe* was significantly lower than in *Ctrl-Read-Low* (*t*_22_ = -2.43, *p* = 0.024) and in *Ctrl-Read-High* (*t*_20_ = -3.83, *p* = 0.001), and marginally lower than in *Dys-Mild* (*t*_24_ = 1.92, *p* = 0.068) and *Ctrl-Age-High* (*t*_20_ = -1.79, *p* = 0.089). Contrastingly, this difference in *Ctrl-Read-High* was marginally higher than in *Ctrl-Age-High* (*t*_22_ = -1.99, *p* = 0.059) and *Ctrl-Read-Low* (*t*_24_ = 1.93, *p* = 0.066).

Linear mixed-effects modeling of syllabic CTS values iteratively revealed a statistically significant effect of noise (χ^2^(2) = 184, *p* < 0.0001), hemisphere (χ^2^(1) = 23.3, *p* < 0.0001) and group (χ^2^(2) = 11.9, *p* = 0.0026), and a significant interaction between hemisphere and group (χ^2^(2) = 6.77, *p* = 0.03). The absence of interaction involving noise indicates that phrasal CTS alteration in dyslexia is not more salient in noisy conditions.

Figure 7 illustrates the effect of noise. The effect was explained by a significant reduction in syllabic CTS in babble noise condition compared with the two other conditions (Noiseless, *t*_77_ = 12.53, *p* < 0.0001; Non-speech noise, *t*_77_ = 12, *p* < 0.0001) for which there was no significant difference (*t*_77_ = 1.70, *p* = 0.09).

**Figure 7.**
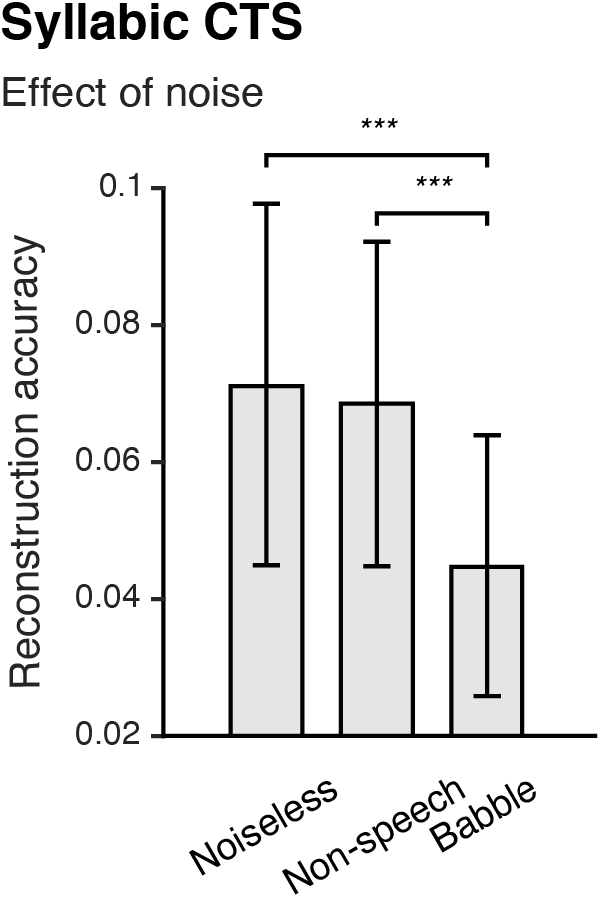
Effect of noise on syllabic CTS. Syllabic CTS was averaged across hemispheres and groups.

The interaction between hemisphere and group was in all points similar to that described in subsection 3.1, for it was not modulated by noise nor subgroup.

## 4. Discussion

This study demonstrates that in French speaking children with dyslexia, (i) syllabic CTS in the right STG is altered in comparison with age-matched controls, but not with reading-level-matched controls, (ii) phrasal/sentential CTS hemispheric lateralization is abnormal in the most severe form of dyslexia, and (iii) phrasal CTS is altered, but only in informational noise, and again, only in comparison with age-matched controls and not with reading-level-matched controls. Overall, these data provide novel insights into the neurobiology of dyslexia that explain the contradictions between previous reports of altered/preserved CTS in dyslexia.

### 4.1. Abnormal syllabic CTS in dyslexia

Our results suggest an abnormal hemispheric lateralization of syllabic CTS in children with dyslexia in comparison with age-matched but not reading-level-matched controls. Syllabic CTS lateralized to the right hemisphere in age-matched controls and showed no sign of lateralization in children with dyslexia and reading-level-matched controls. Accordingly, based on our data, we cannot argue that the alteration in the hemispheric lateralization of syllabic CTS in dyslexia is a core deficit that predates and hampers reading acquisition. The absence of difference between the children with dyslexia and reading-level-matched controls could be due to higher metacognitive abilities in children with dyslexia that would mask a deficit, but it could also be ascribed to reduced reading experience (Goswami, 2015). Indeed, individuals with dyslexia tend to have less scheduled reading time compared with typical readers (Finucci et al., 1985; Sun et al., 2013), and learning to read fosters cerebral development by providing intensive training for sensory (Dehaene et al., 2015) and attentional processes (Goswami, 2015). For example, literacy acquisition improves early visual processes which leads to a reorganization of the ventral occipito-temporal pathway (Szwed et al., 2014) and also modifies phonological coding by strengthening the functional and anatomical link between graphemic and phonemic representations (Dehaene et al., 2015) leading to a reciprocal link between phonology and reading development (Castles and Coltheart, 2004). In the sphere of language, literacy enhances the cerebral activation seen in response to spoken language (Dehaene et al., 2010; Monzalvo and Dehaene-Lambertz, 2013; Nation and Hulme, 2011). Accordingly, reading acquisition improves sensory and attentional brain processes much like maturation does. It is therefore difficult to tell apart the effect of reduced reading experience from that of a developmental disorder itself on sensory or attentional skills. In other words, such deficits can be a cause or a consequence of dyslexia. In our study, the inclusion of —and comparison with— a group of controls matched for reading level was crucial to indicate that the absence of right-hemispheric dominance for syllabic CTS in dyslexia could be attributed to reduced reading experience rather than to a cause of dyslexia.

Syllabic CTS is thought to reflect low-level auditory processing (Molinaro and Lizarazu, 2018) or phonemic processing (Destoky et al., 2020; Di Liberto et al., 2015; Mai et al., 2016). Phonemic processing and hence phonemic awareness have long been posited as causal factors of dyslexia (Tallal, 1980). Phonemic awareness requires a formal teaching, and its acquisition is intertwined with that of reading (Goswami, 2008). Since reading experience influences the development of phonemic awareness, which was here captured by the fidelity of syllabic CTS, it is not surprising that there was a trend for typical readers to show more efficient syllabic tracking than dyslexic readers. In fact, recent evidence suggests that syllabic CTS in dyslexia is similar to that in controls right after speech edges, but decays faster shortly thereafter (Lizarazu et al., 2020).

Furthermore, the development of phonemic awareness is made more difficult for children who learn to read in an orthographically opaque language like French or English (Goswami, 2008; Lallier et al., 2018). Accordingly, the alteration in syllabic CTS in dyslexic readers compared with typical readers of the same age might have been more salient in our study compared with others, because it was conducted on native French speakers.

### 4.2. Altered phrasal CTS lateralization in severe but not mild dyslexia

This study indicates an effect of the severity in reading deficit on the hemispheric dominance of phrasal CTS. Phrasal CTS lateralized to the right hemisphere in children with mild dyslexia while it lateralized to the left hemisphere (though not significantly so) in children with severe dyslexia. Moreover, both control groups showed the same right-hemisphere lateralization for phrasal CTS, indicating that children with severe dyslexia stood out from better readers and younger children with the same reading level with their atypical phrasal CTS lateralization. Their hemispheric lateralization for phrasal CTS is more akin to that previously reported in pre-readers (Ríos-López et al., 2020). The right hemispheric dominance seen in both control groups and in mild dyslexia is in accordance with the asymmetric sampling in time hypothesis, which argues that prosodic and syllabic information of the linguistic signal are preferentially processed in the right hemisphere, while phonemic information (i.e., information at faster rates) would be processed in the left hemisphere or bilaterally (Poeppel, 2003). That hemispheric lateralization was altered in severe dyslexia indicates departure from typical development. In fact, atypical brain hemispheric lateralization during speech processing is commonly reported in dyslexia, and it has been considered as a potential cause of dyslexia (Abrams et al., 2009; Giraud and Poeppel, 2012; Goswami, 2011; Molinaro et al., 2016b). For example, typical readers were reported to present a right-hemispheric dominance for delta and theta band oscillations and a left-hemispheric dominance for gamma band oscillations during an audio-visual perception task, while dyslexic readers of the same age did not present any specific lateralization during this kind of task (Lehongre et al., 2013). This atypical hemispheric lateralization is in line with the temporal sampling framework for developmental dyslexia (Goswami, 2011). This framework argues that the primary neural deficit in dyslexia is impaired phase locking by right-lateralized auditory cortical oscillations at phrasal/syllabic frequencies. Impaired low frequency mechanisms would ultimately hamper the integration between different acoustic features contributing to phonemic perception.

### 4.3. Impact of noise on phrasal CTS in dyslexia

Our results support the well-documented detrimental effect of background informational noise on speech processing in dyslexia (Calcus et al., 2015; Dole et al., 2012; Ziegler et al., 2009). Indeed, phrasal CTS was reduced in dyslexia in comparison with controls in age only in the babble noise condition and not in *noiseless* and *non-speech* noise conditions. This therefore indicates that the alteration in phrasal CTS seen in Spanish or English speaking children with dyslexia (Di Liberto et al., 2018a; Molinaro et al., 2016a; Power et al., 2016a) is also seen in French, but only in challenging listening conditions. It has already been posited that different languages impact differently on some aspects of CTS because of differences in their properties (Lallier et al., 2017). For example, French speakers may tune less strongly their neural oscillations than Spanish or English speakers to slow speech modulations, in particular in the delta (i.e. phrasal) and theta (i.e. syllabic) frequency bands, as a consequence of differences in lexical stress pattern (Lallier et al., 2017). This is because lexical stress in French is totally predictable as it always falls on the last syllable. In contrast, lexical stress in Spanish and English changes depending on the word itself and is used to differentiate between words made of the exact same sequence of phonemes. The perfect predictability of stress in French leads to a “stress deafness” in native speakers (Dupoux et al., 1997), which ultimately results in its underrepresentation of the lexical phonological memory (Dupoux et al., 2010). This French “stress deafness” led us to hypothesize that atypical right-hemisphere neural oscillatory sampling for the low frequencies seen in English and Spanish dyslexic readers would be less severe in French dyslexic readers (Lallier et al., 2017), which is exactly what we have observed. Still, more cross-linguistic studies are warranted to fully determine to which extent language specifics determine the interrelation between reading abilities and the range of language brain functions subtended by CTS.

Notwithstanding the above, the impact of informational noise on phrasal CTS was similar in children with dyslexia compared with their controls in reading level. This, again, suggests that the deficit in phrasal CTS in noise seen in children with dyslexia could be mainly caused by reduced reading experience rather than a causal factor of dyslexia.

Reading experience could impact phrasal CTS in noise through its effect on general lexical knowledge (Destoky et al., 2020). Indeed, reading improves lexical knowledge because new words are more often encountered during reading than listening activities. This effect is commonly called the “Matthew effect” (Morgan et al., 2008). In turn, lexical knowledge influences SiN comprehension (Carroll et al., 2016; Kaandorp et al., 2016; Lewis et al., 2010; Mattys and Wiget, 2011), tentatively through top-down mechanisms that leverage such lexical knowledge to facilitate identification of phonemes by retuning phonemic categories (McClelland et al., 2006). Since the level of phrasal CTS in noise is an electrophysiological correlate of SiN comprehension (Peelle et al., 2013b; Riecke et al., 2018; Vanthornhout et al., 2018), we can surmise the causal chain as follows: reading acquisition develops lexical knowledge which itself boosts the level of phrasal CTS in noise.

### 4.4. Limitations and perspectives

A recurring problem in studies investigating CTS in dyslexia concerns the sample size usually included and the absence of controls in reading level. Indeed, all of these studies included at most 20 dyslexic readers (Di Liberto et al., 2018b; Lizarazu et al., 2021b; Molinaro et al., 2016b; Power et al., 2016b). Also, most of these studies used age-matched controls but no reading-level matched controls (Lizarazu et al., 2021b; Molinaro et al., 2016b; Power et al., 2016b). Our study did slightly better in terms of sample size, and, more importantly, did include appropriate controls in reading level (78 children; 26 children with dyslexia and their controls in age and reading level), making it possible to dissociate potential effects of reading experience from core causes of dyslexia. Despite having included a larger population, we did not identify any specific core alteration in dyslexia that would have been seen in comparison with both control groups. Also, the potential genuine differences we might have missed (false negatives) should have an effect size small enough to dismiss them as core deficits in dyslexia (Friston, 2012). However, this reasoning does not hold for our analysis involving subgroups of participants, where the sample size was substantially reduced; for example, we were left with 10 children with severe dyslexia. Future studies should gather even more participants in order to identify subtler alterations in subgroups (or subtypes) of dyslexia (Saksida et al., 2016).

### 4.5. Conclusion

This study argues that altered CTS in dyslexia is related to reduced reading experience rather than a causal factor of the disorder. Moreover, phrasal tracking showed atypical hemispheric lateralization in children with severe dyslexia but not in those with mild dyslexia. Finally, we demonstrate that phrasal/sentential CTS is not altered in French speaking children dyslexia, in contrast with reports on Spanish or English speaking children. However, the atleration becomes evident in challenging SiN conditions, but again, as a consequence of reduced reading experience.

## Acknowledgments

Florian Destoky, Julie Bertels and Mathieu Bourguignon have been supported by the program Attract of Innoviris (grants 2015-BB2B-10 and 2019-BFB-110). Julie Bertels has been supported by a research grant from the Fonds de Soutien Marguerite-Marie Delacroix (Brussels, Belgium). Xavier De Tiège is Post-doctorate Clinical Master Specialist at the Fonds de la Recherche Scientifique (F.R.S.-FNRS, Brussels, Belgium). Mathieu Bourguignon has been supported by the Marie Skłodowska-Curie Action of the European Commission (grant 743562).

The MEG project at the CUB Hôpital Erasme and this study were financially supported by the Fonds Erasme (Research convention “Les Voies du Savoir”, Brussels, Belgium).

The PET-MR project at the CUB Hôpital Erasme is supported by the Association Vinçotte Nuclear (AVN, Brussels, Belgium).

## Notes

**Conflicts of interest** None of the authors disclose any potential conflict of interest.

### Competing Interest Statement

The authors have declared no competing interest.

## References

1. Abrams DA, Nicol T, Zecker S, Kraus N. 2009. Abnormal cortical processing of the syllable rate of speech in poor readers. J Neurosci 29:7686–7693.

2. Ahissar E, Nagarajan S, Ahissar M, Protopapas A, Mahncke H, Merzenich MM. 2001. Speech comprehension is correlated with temporal response patterns recorded from auditory cortex. Proc Natl Acad Sci U S A 98:13367–13372.

3. Ashburner J, Friston KJ. 1999. Nonlinear spatial normalization using basis functions. Hum Brain Mapp 7:254–266.

4. Ashburner J, Neelin P, Collins DL, Evans A, Friston K. 1997. Incorporating prior knowledge into image registration. Neuroimage 6:344–352.

5. Bates D, Mächler M, Bolker B, Walker S. 2015. Fitting Linear Mixed-Effects Models Using lme4. Journal of Statistical Software. doi:10.18637/jss.v067.i01

6. Biesmans W, Das N, Francart T, Bertrand A. 2017. Auditory-Inspired Speech Envelope Extraction Methods for Improved EEG-Based Auditory Attention Detection in a Cocktail Party Scenario. IEEE Transactions on Neural Systems and Rehabilitation Engineering. doi:10.1109/tnsre.2016.2571900

7. Bonte ML, Poelmans H, Blomert L. 2007. Deviant neurophysiological responses to phonological regularities in speech in dyslexic children. Neuropsychologia 45:1427–1437.

8. Bourguignon M, De Tiège X, Op de Beeck M, Ligot N, Paquier P, Van Bogaert P, Goldman S, Hari R, Jousmäki V. 2013. The pace of prosodic phrasing couples the listener’s cortex to the reader’s voice. Human Brain Mapping. doi:10.1002/hbm.21442

9. Bourguignon M, Jousmäki V, Op de Beeck M, Van Bogaert P, Goldman S, De Tiège X. 2012. Neuronal network coherent with hand kinematics during fast repetitive hand movements. NeuroImage. doi:10.1016/j.neuroimage.2011.09.022

10. Bourguignon M, Molinaro N, Wens V. 2018. Contrasting functional imaging parametric maps: The mislocation problem and alternative solutions. NeuroImage. doi:10.1016/j.neuroimage.2017.12.033

11. Calcus A, Colin C, Deltenre P, Kolinsky R. 2015. Informational masking of speech in dyslexic children. The Journal of the Acoustical Society of America. doi:10.1121/1.4922012

12. Carreiras M, Seghier ML, Baquero S, Estévez A, Lozano A, Devlin JT, Price CJ. 2009. An anatomical signature for literacy. Nature 461:983–986.

13. Carroll R, Warzybok A, Kollmeier B, Ruigendijk E. 2016. Age-Related Differences in Lexical Access Relate to Speech Recognition in Noise. Front Psychol 7:990.

14. Castles A, Coltheart M. 2004. Is there a causal link from phonological awareness to success in learning to read? Cognition. doi:10.1016/s0010-0277(03)00164-1

15. Crosse MJ, Di Liberto GM, Bednar A, Lalor EC. 2016. The Multivariate Temporal Response Function (mTRF) Toolbox: A MATLAB Toolbox for Relating Neural Signals to Continuous Stimuli. Front Hum Neurosci 10:604.

16. Dale AM, Sereno MI. 1993. Improved Localizadon of Cortical Activity by Combining EEG and MEG with MRI Cortical Surface Reconstruction: A Linear Approach. Journal of Cognitive Neuroscience. doi:10.1162/jocn.1993.5.2.162

17. Dehaene S, Cohen L, Morais J, Kolinsky R. 2015. Illiterate to literate: behavioural and cerebral changes induced by reading acquisition. Nat Rev Neurosci 16:234–244.

18. Dehaene S, Pegado F, Braga LW, Ventura P, Nunes Filho G, Jobert A, Dehaene-Lambertz G, Kolinsky R, Morais J, Cohen L. 2010. How learning to read changes the cortical networks for vision and language. Science 330:1359–1364.

19. Demanez L, Dony-Closon B, Lhonneux-Ledoux E, Demanez JP. 2003. Central auditory processing assessment: a French-speaking battery. Acta Otorhinolaryngol Belg 57:275–290.

20. Destoky F, Bertels J, Niesen M, Wens V, Vander Ghinst M, Leybaert J, Lallier M, Ince RAA, Gross J, De Tiège X, Bourguignon M. 2020. Cortical tracking of speech in noise accounts for reading strategies in children. PLoS Biol 18:e3000840.

21. Destoky F, Philippe M, Bertels J, Verhasselt M, Coquelet N, Vander Ghinst M, Wens V, De Tiège X, Bourguignon M. 2019a. Comparing the potential of MEG and EEG to uncover brain tracking of speech temporal envelope. Neuroimage 184:201–213.

22. Destoky F, Philippe M, Bertels J, Verhasselt M, Coquelet N, Vander Ghinst M, Wens V, De Tiège X, Bourguignon M. 2019b. Comparing the potential of MEG and EEG to uncover brain tracking of speech temporal envelope. Neuroimage 184:201–213.

23. De Tiège X, Op de Beeck M, Funke M, Legros B, Parkkonen L, Goldman S, Van Bogaert P. 2008. Recording epileptic activity with MEG in a light-weight magnetic shield. Epilepsy Research. doi:10.1016/j.eplepsyres.2008.08.011

24. Di Liberto GM, O’Sullivan JA, Lalor EC. 2015. Low-Frequency Cortical Entrainment to Speech Reflects Phoneme-Level Processing. Curr Biol 25:2457–2465.

25. Di Liberto GM, Peter V, Kalashnikova M, Goswami U, Burnham D, Lalor EC. 2018a. Atypical cortical entrainment to speech in the right hemisphere underpins phonemic deficits in dyslexia. Neuroimage 175:70–79.

26. Di Liberto GM, Peter V, Kalashnikova M, Goswami U, Burnham D, Lalor EC. 2018b. Atypical cortical entrainment to speech in the right hemisphere underpins phonemic deficits in dyslexia. Neuroimage 175:70–79.

27. Ding N, Melloni L, Zhang H, Tian X, Poeppel D. 2016. Cortical tracking of hierarchical linguistic structures in connected speech. Nat Neurosci 19:158–164.

28. Ding N, Simon JZ. 2014. Cortical entrainment to continuous speech: functional roles and interpretations. Front Hum Neurosci 8:311.

29. Dole M, Hoen M, Meunier F. 2012. Speech-in-noise perception deficit in adults with dyslexia: Effects of background type and listening configuration. Neuropsychologia. doi:10.1016/j.neuropsychologia.2012.03.007

30. Dupoux E, Pallier C, Sebastian N, Mehler J. 1997. A Destressing “Deafness” in French? Journal of Memory and Language. doi:10.1006/jmla.1996.2500

31. Dupoux E, Peperkamp S, Sebastián-Gallés N. 2010. Limits on bilingualism revisited: stress “deafness” in simultaneous French-Spanish bilinguals. Cognition 114:266–275.

32. Finucci JM, Gottfredson LS, Childs B. 1985. A follow-up study of dyslexic boys. Annals of Dyslexia. doi:10.1007/bf02659183

33. Friston K. 2012. Ten ironic rules for non-statistical reviewers. Neuroimage 61:1300–1310.

34. Giraud A-L, Poeppel D. 2012. Cortical oscillations and speech processing: emerging computational principles and operations. Nat Neurosci 15:511–517.

35. Golumbic EZ, Zion Golumbic E, Cogan GB, Schroeder CE, Poeppel D. 2013. Visual Input Enhances Selective Speech Envelope Tracking in Auditory Cortex at a “Cocktail Party.” Journal of Neuroscience. doi:10.1523/jneurosci.3675-12.2013

36. Goswami U. 2015. Sensory theories of developmental dyslexia: three challenges for research. Nat Rev Neurosci 16:43–54.

37. Goswami U. 2011. A temporal sampling framework for developmental dyslexia. Trends Cogn Sci 15:3–10.

38. Goswami U. 2008. Reading, complexity and the brain. Lit Discuss 42:67–74.

39. Gramfort A, Luessi M, Larson E, Engemann DA, Strohmeier D, Brodbeck C, Parkkonen L, Hämäläinen MS. 2014. MNE software for processing MEG and EEG data. Neuroimage 86:446–460.

40. Gross J, Hoogenboom N, Thut G, Schyns P, Panzeri S, Belin P, Garrod S. 2013. Speech rhythms and multiplexed oscillatory sensory coding in the human brain. PLoS Biol 11:e1001752.

41. Hämäläinen JA, Lohvansuu K, Ervast L, Leppänen PHT. 2015. Event-related potentials to tones show differences between children with multiple risk factors for dyslexia and control children before the onset of formal reading instruction. Int J Psychophysiol 95:101–112.

42. Hämäläinen MS, Ilmoniemi RJ. 1994. Interpreting magnetic fields of the brain: minimum norm estimates. Med Biol Eng Comput 32:35–42.

43. Jacquier-Roux M, Valdois S, Zorman M. 2002. Odédys: outil de dépistage des dyslexies.

44. Kaandorp MW, De Groot AMB, Festen JM, Smits C, Goverts ST. 2016. The influence of lexical-access ability and vocabulary knowledge on measures of speech recognition in noise. Int J Audiol 55:157–167.

45. Krashen S. 1999. Training in Phonemic Awareness: Greater on Tests of Phonemic Awareness. Perceptual and Motor Skills. doi:10.2466/pms.1999.89.2.412

46. Lachmann T, Weis T. 2018. Reading and Dyslexia: From Basic Functions to Higher Order Cognition. Springer.

47. Lallier M, Lizarazu M, Molinaro N, Bourguignon M, Ríos-López P, Carreiras M. 2018. From Auditory Rhythm Processing to Grapheme-to-Phoneme Conversion: How Neural Oscillations Can Shed Light on Developmental Dyslexia. Literacy Studies. doi:10.1007/978-3-319-90805-2_8

48. Lallier M, Molinaro N, Lizarazu M, Bourguignon M, Carreiras M. 2017. Amodal Atypical Neural Oscillatory Activity in Dyslexia. Clinical Psychological Science. doi:10.1177/2167702616670119

49. Lefavrais P. 2005. Manuel du test de l’alouette.

50. Lehongre K, Morillon B, Giraud A-L, Ramus F. 2013. Impaired auditory sampling in dyslexia: further evidence from combined fMRI and EEG. Front Hum Neurosci 7:454.

51. Leong V, Goswami U. 2014. Impaired extraction of speech rhythm from temporal modulation patterns in speech in developmental dyslexia. Front Hum Neurosci 8:96.

52. Leppänen PHT, Hämäläinen JA, Guttorm TK, Eklund KM, Salminen H, Tanskanen A, Torppa M, Puolakanaho A, Richardson U, Pennala R, Lyytinen H. 2012. Infant brain responses associated with reading-related skills before school and at school age. Neurophysiol Clin 42:35–41.

53. Lewis D, Hoover B, Choi S, Stelmachowicz P. 2010. Relationship between speech perception in noise and phonological awareness skills for children with normal hearing. Ear Hear 31:761–768.

54. Lizarazu M, di Covella LS, van Wassenhove V, Rivière D, Mizzi R, Lehongre K, Hertz-Pannier L, Ramus F. 2021a. Neural entrainment to speech and nonspeech in dyslexia: conceptual replication and extension of previous investigations. Cortex. doi:10.1016/j.cortex.2020.12.024

55. Lizarazu M, di Covella LS, van Wassenhove V, Rivière D, Mizzi R, Lehongre K, Hertz-Pannier L, Ramus F. 2021b. Neural entrainment to speech and nonspeech in dyslexia: conceptual replication and extension of previous investigations. Cortex. doi:10.1016/j.cortex.2020.12.024

56. Lizarazu M, Lallier M, Bourguignon M, Carreiras M, Molinaro N. 2020. Impaired neural response to speech edges in dyslexia. Cortex 135:207–218.

57. Luo H, Poeppel D. 2007. Phase patterns of neuronal responses reliably discriminate speech in human auditory cortex. Neuron 54:1001–1010.

58. Lyon GR, Reid Lyon G, Shaywitz SE, Shaywitz BA. 2003. A definition of dyslexia. Annals of Dyslexia. doi:10.1007/s11881-003-0001-9

59. Mai G, Minett JW, Wang WS-Y. 2016. Delta, theta, beta, and gamma brain oscillations index levels of auditory sentence processing. Neuroimage 133:516–528.

60. Mattys SL, Wiget L. 2011. Effects of cognitive load on speech recognition. Journal of Memory and Language. doi:10.1016/j.jml.2011.04.004

61. McClelland JL, Mirman D, Holt LL. 2006. Are there interactive processes in speech perception? Trends Cogn Sci 10:363–369.

62. Meyer L, Gumbert M. 2018. Synchronization of Electrophysiological Responses with Speech Benefits Syntactic Information Processing. J Cogn Neurosci 30:1066–1074.

63. Meyer L, Henry MJ, Gaston P, Schmuck N, Friederici AD. 2017. Linguistic Bias Modulates Interpretation of Speech via Neural Delta-Band Oscillations. Cereb Cortex 27:4293–4302.

64. Molinaro N, Lizarazu M. 2018. Delta(but not theta)-band cortical entrainment involves speech-specific processing. Eur J Neurosci 48:2642–2650.

65. Molinaro N, Lizarazu M, Lallier M, Bourguignon M, Carreiras M. 2016a. Out-of-synchrony speech entrainment in developmental dyslexia. Hum Brain Mapp 37:2767–2783.

66. Molinaro N, Lizarazu M, Lallier M, Bourguignon M, Carreiras M. 2016b. Out-of-synchrony speech entrainment in developmental dyslexia. Hum Brain Mapp 37:2767–2783.

67. Monzalvo K, Dehaene-Lambertz G. 2013. How reading acquisition changes children’s spoken language network. Brain Lang 127:356–365.

68. Morgan PL, Farkas G, Hibel J. 2008. Matthew Effects for Whom? Learn Disabil Q 31:187–198.

69. Myronenko A, Song X. 2010. Point set registration: coherent point drift. IEEE Trans Pattern Anal Mach Intell 32:2262–2275.

70. Nation K, Hulme C. 2011. Learning to read changes children’s phonological skills: evidence from a latent variable longitudinal study of reading and nonword repetition. Developmental Science. doi:10.1111/j.1467-7687.2010.01008.x

71. Nichols TE, Holmes AP. 2002. Nonparametric permutation tests for functional neuroimaging: a primer with examples. Hum Brain Mapp 15:1–25.

72. Olson RK, Wise B, Ring J, Johnson M. 1997. Computer-Based Remedial Training in Phoneme Awareness and Phonological Decoding: Effects on the Posttraining Development of Word Recognition. Scientific Studies of Reading. doi:10.1207/s1532799xssr0103_4

73. Oostenveld R, Fries P, Maris E, Schoffelen J-M. 2011. FieldTrip: Open source software for advanced analysis of MEG, EEG, and invasive electrophysiological data. Comput Intell Neurosci 2011:156869.

74. Pape-Neumann J, van Ermingen-Marbach M, Grande M, Willmes K, Heim S. 2015. The role of phonological awareness in treatments of dyslexic primary school children. Acta Neurobiol Exp 75:80–106.

75. Park H, Ince RAA, Schyns PG, Thut G, Gross J. 2018. Representational interactions during audiovisual speech entrainment: Redundancy in left posterior superior temporal gyrus and synergy in left motor cortex. PLoS Biol 16:e2006558.

76. Park H, Kayser C, Thut G, Gross J. 2016. Lip movements entrain the observers’ low-frequency brain oscillations to facilitate speech intelligibility. eLife. doi:10.7554/elife.14521

77. Paz-Alonso PM, Oliver M, Lerma-Usabiaga G, Caballero-Gaudes C, Quiñones I, Suárez-Coalla P, Duñabeitia JA, Cuetos F, Carreiras M. 2018. Neural correlates of phonological, orthographic and semantic reading processing in dyslexia. Neuroimage Clin 20:433–447.

78. Peelle JE, Gross J, Davis MH. 2013a. Phase-locked responses to speech in human auditory cortex are enhanced during comprehension. Cereb Cortex 23:1378–1387.

79. Peelle JE, Gross J, Davis MH. 2013b. Phase-locked responses to speech in human auditory cortex are enhanced during comprehension. Cereb Cortex 23:1378–1387.

80. Poeppel D. 2003. The analysis of speech in different temporal integration windows: cerebral lateralization as “asymmetric sampling in time.” Speech Communication. doi:10.1016/s0167-6393(02)00107-3

81. Pollack I. 1975. Auditory informational masking. The Journal of the Acoustical Society of America. doi:10.1121/1.1995329

82. Power AJ, Colling LJ, Mead N, Barnes L, Goswami U. 2016a. Neural encoding of the speech envelope by children with developmental dyslexia. Brain and Language. doi:10.1016/j.bandl.2016.06.006

83. Power AJ, Colling LJ, Mead N, Barnes L, Goswami U. 2016b. Neural encoding of the speech envelope by children with developmental dyslexia. Brain and Language. doi:10.1016/j.bandl.2016.06.006

84. Reuter M, Schmansky NJ, Rosas HD, Fischl B. 2012. Within-subject template estimation for unbiased longitudinal image analysis. Neuroimage 61:1402–1418.

85. Riecke L, Formisano E, Sorger B, Başkent D, Gaudrain E. 2018. Neural Entrainment to Speech Modulates Speech Intelligibility. Curr Biol 28:161–169.e5.

86. Ríos-López P, Molinaro N, Bourguignon M, Lallier M. 2020. Development of neural oscillatory activity in response to speech in children from 4 to 6 years old. Developmental Science. doi:10.1111/desc.12947

87. Saksida A, Iannuzzi S, Bogliotti C, Chaix Y, Démonet J-F, Bricout L, Billard C, Nguyen-Morel M-A, Le Heuzey M-F, Soares-Boucaud I, George F, Ziegler JC, Ramus F. 2016. Phonological skills, visual attention span, and visual stress in developmental dyslexia. Dev Psychol 52:1503–1516.

88. Sun Z, Zou L, Zhang J, Mo S, Shao S, Zhong R, Ke J, Lu X, Miao X, Song R. 2013. Prevalence and Associated Risk Factors of Dyslexic Children in a Middle-Sized City of China: A Cross-Sectional Study. PLoS ONE. doi:10.1371/journal.pone.0056688

89. Szwed M, Qiao E, Jobert A, Dehaene S, Cohen L. 2014. Effects of literacy in early visual and occipitotemporal areas of Chinese and French readers. J Cogn Neurosci 26:459–475.

90. Tallal P. 1980. Auditory temporal perception, phonics, and reading disabilities in children. Brain Lang 9:182–198.

91. Taulu S, Simola J. 2006. Spatiotemporal signal space separation method for rejecting nearby interference in MEG measurements. Phys Med Biol 51:1759–1768.

92. Taulu S, Simola J, Kajola M. 2005. Applications of the signal space separation method. IEEE Transactions on Signal Processing. doi:10.1109/tsp.2005.853302

93. Team RC, Others. 2013. R: A language and environment for statistical computing.

94. Vander Ghinst M, Bourguignon M, Niesen M, Wens V, Hassid S, Choufani G, Jousmäki V, Hari R, Goldman S, De Tiège X. 2019a. Cortical Tracking of Speech-in-Noise Develops from Childhood to Adulthood. J Neurosci 39:2938–2950.

95. Vander Ghinst M, Bourguignon M, Niesen M, Wens V, Hassid S, Choufani G, Jousmäki V, Hari R, Goldman S, De Tiège X. 2019b. Cortical Tracking of Speech-in-Noise Develops from Childhood to Adulthood. J Neurosci 39:2938–2950.

96. Vander Ghinst M, Bourguignon M, Op de Beeck M, Wens V, Marty B, Hassid S, Choufani G, Jousmäki V, Hari R, Van Bogaert P, Goldman S, De Tiège X. 2016. Left Superior Temporal Gyrus Is Coupled to Attended Speech in a Cocktail-Party Auditory Scene. J Neurosci 36:1596–1606.

97. Vanthornhout J, Decruy L, Wouters J, Simon JZ, Francart T. 2018. Speech intelligibility predicted from neural entrainment of the speech envelope. Journal of the Association for Research in Otolaryngology. doi:10.1101/246660

98. Wens V, Marty B, Mary A, Bourguignon M, Op de Beeck M, Goldman S, Van Bogaert P, Peigneux P, De Tiège X. 2015. A geometric correction scheme for spatial leakage effects in MEG/EEG seed-based functional connectivity mapping. Hum Brain Mapp 36:4604–4621.

99. Ziegler JC, Pech-Georgel C, George F, Lorenzi C. 2009. Speech-perception-in-noise deficits in dyslexia. Developmental Science. doi:10.1111/j.1467-7687.2009.00817.x

